# piRNAdb: A piwi-interacting RNA database

**DOI:** 10.1101/2021.09.21.461238

**Authors:** Ricardo Piuco, Pedro A. F. Galante

## Abstract

**Motivation:** Found in several metazoan species, piwi-interacting RNAs (piRNAs) regulate the expression of a wide variety of transposable elements and genes associated with cellular development, differentiation, and growth. Despite their importance, piRNAs are not well known and are still underexplored. To facilitate piRNA research, a comprehensive and easy-to-use piRNA database is still needed.

**Results:** Here, we present piRNAdb, an integrative and user-friendly database designed to encompass several aspects of piRNAs. We selected piRNAs from four reliable human RNA-sequencing datasets to start our database. After data processing, we displayed these sequences, their genomic location, clustering information, putative targets on known genes and transposable elements, as well as direct links to other databases. In this first release, piRNAdb catalogues 27,329 piRNAs, as well as 23,380 genes that are putative targets and 47,060 associated gene ontology terms, both of which are organized and linked to each respective piRNA. Finally, to improve information exchange and increase the confidence of sequences, a feedback system is provided to users of piRNAdb.

**Conclusion:** The inclusion of new features to facilitate piRNA analyses, data visualization, and integration is the major pillar of piRNAdb. Our main goal was to make this database an easy interface between the data and the user. We believe that this web tool achieves this objective by providing a streamlined and well-organized data repository for piRNAs and that it will be extremely useful to those already studying piRNAs and to the broader community.

**Availability:** piRNAdb is available freely and is compatible with smartphones and tablets: https://www.pirnadb.org/

## Introduction

*Piwi-interacting* RNA (piRNA) is a class of small non-coding RNAs that range from 26 to 31 nucleotides (1). piRNAs are involved in the regulation of several biological processes, including i) control of transposable element (TE) expression (2); ii) renewal and differentiation of stem cells (3); iii) animal development (4); and iv) invasion, differentiation, and growth of breast tumors, gastric tumors, and hepatocellular carcinomas (5).

piRNAs are found in almost all metazoans with surprisingly diversity, even for closely related species (6). Most piRNAs originate from large genomic regions, named clusters (7), that have a high density of piRNA sequences. These clusters are known ‘traps’ for TEs; after a truncated form of this element is inserted into a piRNA cluster, piRNA transcription can repress the parental element. This event increases the repertory and diversity of piRNAs that control the expression of cell molecules (7). The bound piRNA-PIWI protein acts inside the cell nucleus, recruiting methylation or breaking the non-processed mRNAs of their targets (8). As an increasing number of diverse piRNAs are identified, a comprehensive and user-friendly database is needed. Although some piRNA databases are available, none of them provide a complete, updated, integrated, and user-friendly set of information about piRNAs. For example, piRNABank (9) stores only data from NCBI and does not include additional information relevant to these short RNAs. piRBase (10) has the greatest number of piRNA sequences; however, it also presents suspicious piRNAs that lack alignment to the human genome.

Here, we aimed to develop a comprehensive database of piRNAs. Our resulting piwi-interacting RNA database (piRNAdb) groups all known piRNA sequences in humans and includes information about alignments to the genome, putative targets of piRNAs, clusters of piRNAs, gene ontology classification for target genes, and hotlinks to other databases. Importantly, our freely available web-based tool uses modern programming patterns and architectures, allowing easy, intuitive, and responsive browsing with all devices, including mobiles.

### Data Implementation

We created and implemented a MySQL (www.mysql.com) relational database to store all of the data presented in the piRNAdb web interface. We developed the back-end interface using the Model-View-Controller architecture in PHP language (http://php.net) due to its ideal data separation and system developer group workability. We developed our front-end interface using HyperText Markup Language (HTML) 5, Asynchronous JavaScript, and Cascading Style Sheets 3.

To assign specific pages and information sections more relevance, we incorporated search engine optimization techniques and semantic HTML in order to improve the ranking on search engines. In an effort to make data retrieval faster and more accessible, we developed piRNAdb following responsive rules, which are specific instructions given to different devices to display the website content in correct proportions. As a result, our database is functional on the majority of modern browsers, tablets, and smartphones (mobiles).

### Data Sources

Our complete database includes all piRNA sequences already reported in humans according to datasets from (1,11–13). We built a PHP script to process these piRNA data and upload the information to piRNAdb (see next section for details). Of the 34,322 piRNA sequences, 1,496 were found to be duplicated in two or more datasets; therefore, we merged these sequences. Nevertheless, we kept references to all original data sources, even for merged piRNAs. We downloaded the human genome (versions: hg19 and hg38) and transcriptome (RELEASE 84) from the University of California, Santa Cruz (UCSC) genome browser (http://genome.ucsc.edu) and ENSEMBL (http://www.ensembl.org), respectively.

### Mapping piRNAs

For gaining information about piRNA genome location, we first aligned the total set of 32,826 (34,322 – 1,496) piRNAs against the human genome (hg19 and hg38). We used the Burrows-Wheeler Aligner (parameters: -n 0 -t 8) (14), which does not allow mismatches or gaps, and kept only full-length sequence alignments against the genome. Using this approach, we identified approximately 1,600,000 alignment positions (789,637 on hg19 and 794,576 on hg38).

Human chromosomes 15 (37,938 alignments), 17 (37,174 alignments) and 19 (34,183 alignments) had approximately the same amount of alignments as chromosome 4 (38,812 alignments), despite the fact that they (15, 17, and 19) are shorter in length than chromosome 4. Curiously, chromosomes 19 and 17 had the highest density of coding genes in our genome (ENSEMBL RELEASE 87). Likewise, chromosome 15 had the highest density of small nucleolar RNAs among all human chromosomes (ENSEMBL RELEASE 67).

Next, we developed an in-house pipeline to process and upload the alignments to piRNAdb. We found that 26,879 (82%) of all piRNAs perfectly matched to the human genome. In total, 437 piRNAs (1.34%) presented one or two mismatches within polymorphic sites (based on NCBI’s dbSNP, version 150). The remaining 5,510 piRNAs presented one or two mismatches at other genomic regions (hg19 or hg38). We only kept piRNAs with perfect matches against the human genome in piRNAdb. However, the full list of piRNAs can be found and downloaded directly on the piRNAdb website.

### Clusters

We next determined the piRNAs clusters using a similar approach as (15). In brief, we searched for 20kb genomic windows (with a 1kb step) that contained at least seven piRNA alignments. We grouped piRNAs matching this cut-off into clusters. In this step, we used only those piRNAs with unique alignment to the genome. In total, we detected 217 clusters using this methodology. The number of clusters found did not correlate with the chromosome size; for example, chromosome 15 had more clusters (28) than chromosome 1 (10), despite being greater in length and having a larger number of piRNAs. Chromosome 17 and 19, which have the highest number of piRNAs, showed a similar number of clusters as other chromosomes.

### Targets

The last step of our data processing pipeline was the detection of piRNA targets. We searched for piRNA targets within the lists of all known TEs and coding and non-coding genes. To be considered a target, we required the genomic coordinates to overlap between a piRNA and at least one transcript from a gene. We allowed overlapping on both exonic and intronic regions, as piRNAs are known to bind non-processed mRNA (16).

We found a total of 262,458 alignments from 18,799 unique piRNAs in which the genomic coordinates overlapped with transcripts; 14,801 were located on protein-coding regions and 6,163 on non-coding regions. This data suggests that, similar to microRNAs (miRNAs), a piRNA may have several genes as targets and a gene may be targeted by several piRNAs (Supplementary Figure 1 and Supplementary Figure 2). For example, DNM1P47 is a potential target for 251 piRNAs, and piRNA hsa-piR-20792 has targets in 26,533 genes. Interesting, among the hsa-piR-13284 targets, we found genes already described as being expressed in few tissues or being broadly expressed. For example, CNTNAP2 is known to be expressed during nervous system development (Poot, 2015) and EIF4G3 is a validated miRNA target that is involved in mRNA cap recognition and transport to ribosomes (17). Therefore, our results show that piRNAs have targets not only in TEs (see below), but also within coding and non-coding genes, including those widely expressed and those with known functions in specific organs/tissues.

We applied a similar methodology to find piRNA targets within TEs. We found 228,869 piRNAs alignments (representing 3,680 unique piRNAs) overlapping with TEs. Interesting, 1,076 (4%) piRNAs had targets in both protein-coding genes and TEs elements, whereas other piRNAs (1,400; 5.20%) bound only to TEs. Coding/non-coding gene and TE targets for piRNAs are displayed in the same section (piRNA Alignments) on the piRNAdb web tool.

We next aimed to identify ontology terms related to piRNA targets. Approximately 47,000 gene ontology terms (18) were available to the piRNA targets. The top three most recurrent biological process terms were (1) transcription, (2) regulation of transcription, and (3) signal transduction (1,366, 992, and 820 genes annotated, respectively). The three most frequent molecular function terms were (1) protein binding, (2) metal ion binding, and (3) ATP binding (7,135, 1,858, and 1,292 genes annotated, respectively). These findings are shown in Supplementary Figure 3. To the best of our knowledge, piRNAdb is the first piRNA database to display gene ontology information. We believe that this information will be advantageous to users in performing networking and integrative analyses using piRNAs’ targets.

### Database Query and Interface

Currently, the field lacks a standard nomenclature for piRNAs. In order to contribute to a standardization, we named piRNAs following a similar pattern used for miRNA nomenclature. We assigned the piRNAs a unique code formed by the initial related to the organism (hsa, for human), the word “piR” (meaning piRNA), and a numerical code (e.g., resulting in hsa-piR-1 or hsa-piR-5). The identified clusters also received similar codes formed by the initial organism, the word “clu”, and a numerical code (e.g., hsa-clu-7 and hsa-clu-17).

We also developed a tool to link piRNAs named in other databases to our system and vice versa. In the “Cross Codes” section, the user is able to place a piRNA named in another database; our system then finds and displays the access code in piRNAdb. Beyond the piRNA name, retrieving information can be performed using the chromosome position, cluster name, ENSEMBL code ID, or gene name of a desired target. For example, if the user searches for a piRNA target (Figure 1a), a list of piRNAs (if they exist) will be retrieved (Figure 1b). By clicking on a piRNA, a webpage displays further information about the selected piRNA, such as its nucleotide sequence (Figure 1c), alignments to the genome, and information about genes and transposable elements that may be associated with each alignment (Figure 1d). In order to better integrate our data with other databases, we included hotlinks on piRNAdb. For instance, clicking on one of piRNAs alignments will redirect the user to the UCSC genome browser page, showing the respective piRNA genomic coordinates and other information.

**Figure 1.**
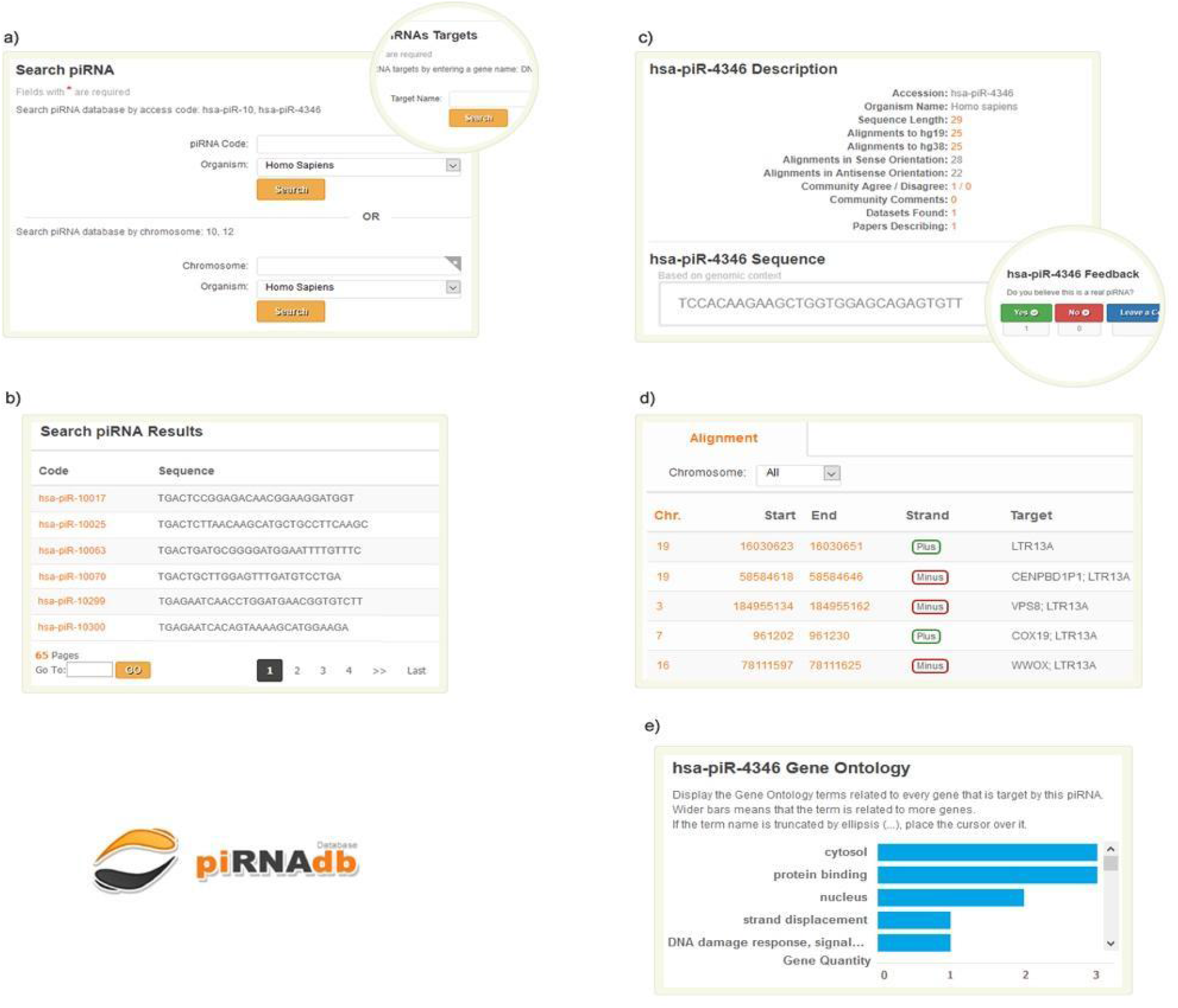
Schematic Search process and information accession about a specific piRNA (has-piR-4346). a) Search form providing some options. Circle on top right displays the search by official gene name to find piRNAs that possibly target it. b) piRNA list that fulfills the term on the search step. c) Information page for a specific piRNA with feedback system on the bottom right circle. d) Section listing all the alignments to the genome (version hg38), putative gene and transposable elements associated with the hsa-piRNA-4346. e) Gene Ontology terms associated with the gene that are (putative) targets of hsa-piR-4346.

Also unique to this database, piRNAdb provides an interactive system where users may give feedback about the reliability of a piRNA (Figure 1c). At least for now, we will analyze and curate all feedback. It is also possible for the user to be directed to Wikipedia (www.wikipedia.org) to describe the piRNA there as well. These features will enable faster communication among researchers and enthusiasts of discoveries related to piRNAs.

Aiming for an easier visualization of the targets related to piRNAs for the user and search engines, we developed a section below the alignments to show all the targets for a specific piRNA in the tag cloud format. This feature shows the most recurrent genes in a larger font size and stronger color to highlight them to the users. Associated gene ontology terms to these gene targets are displayed below (Figure 1e); information about which genes have each term and the frequency of occurrence is available as well.

At the end of the page, the “Datasets” section displays information about the methodology and tissues studied for the piRNA characterization. Finally, a list of all the references linked to each specific piRNA is shown. All raw data displayed in our web interface can be downloaded (“Downloads” page) in various formats. A frequently asked questions and help section (FAQ/Help page) provides information to the users about using the web interface, the data version, and potential browsing problems. As this is the first piRNAdb version (RELEASE 1), it only provides information about humans. We anticipate that in the next release, piRNAdb will include piRNAs and their related information for an additional organism as well.

## Conclusion

Our system provides a powerful and user-friendly web tool for studying piRNAs. piRNAdb uses modern programming architecture and tools, as well as techniques to ensure scalability, usability, and better visualization for users. Our web tool displays only relevant information about piRNAs, within a streamlined and well-organized interface that is compatible with almost all devices, including mobiles. We also provide information that is currently lacking in other piRNA databases, such as gene ontology terms. In addition, as we performed a strict selection of the piRNAs stored on our database, our system is more robust and reliable for use by researchers.

In conclusion, piRNAdb provides reliable information about piRNAs and several downstream analyses to help facilitate piRNA studies. Our findings on the amount of alignments and putative gene targets associated with gene ontologies provide unique information about our piRNAs. In the future, strong candidates could be further explored or identified by utilizing the piRNAdb system.

## Acknowledgements

We thank members of the Galante lab for suggestions and fruitful discussions. R.P. is supported by a Coordenação de Aperfeiçoamento de Pessoal de Nível Superior fellowship. This work was in part funded by the Fundação de Amparo à Pesquisa do Estado de São Paulo (2012/24731-1 to P.A.F.G.) and CNPq (to P.A.F.G.).

